# Rational design of hidden thermodynamic driving through DNA mismatch repair

**DOI:** 10.1101/426668

**Authors:** Natalie E. C. Haley, Thomas E. Ouldridge, Alessandro Geraldini, Ard A. Louis, Jonathan Bath, Andrew J. Turberfield

## Abstract

Recent years have seen great advances in the development of synthetic self-assembling molecular systems. Designing out-of-equilibrium architectures, however, requires a more subtle control over the thermodynamics and kinetics of reactions. We propose a new mechanism for enhancing thermodynamic drive of DNA strand displacement reactions whilst barely perturbing forward reaction rates - introducing mismatches in an internal location within the initial duplex. Through a combination of experiment and simulation, we demonstrate that displacement rates are strongly sensitive to mismatch location and can be tuned by rational design. By placing mismatches away from duplex ends, the thermodynamic drive for a strand-displacement reaction can be varied without significantly affecting the forward reaction rate. This hidden thermodynamic driving motif is ideal for the engineering of nonequilibrium systems that rely on catalytic control and must be robust to leak reactions.

---

One of the signature features of living systems is that they operate continuously, expending free energy, rather than relaxing to equilibrium. Key molecular components are not consumed by these reactions but recovered – they act as catalysts.^1^ Examples include metabolic enzymes, signal-processing kinases, molecular motors, polymerases and even nucleic acids undergoing replication, transcription and translation.^2^

Designing synthetic analogues of such complex molecular systems is a key goal of nanotechnology. This task is complicated by the requirements that reactions must be thermodynamically favourable to proceed while stable equilibrium states that lock up the key catalytic components, and accidental leak reactions, must be avoided. Overall reaction thermodynamics and kinetics must therefore be carefully tuned.

Due to its predictable base-pairing interactions, DNA has proved to be a remarkable material for the construction of static nanoscale structures^3–7^ and systems that perform single-shot computations.^8,9^ Continuouly operating dynamic systems, including reaction network architectures^9-11^ and synthetic molecular machinery,^12-14^ have also been developed, but the state of the art is far from the power and flexibility of natural analogs.

Toehold-mediated strand displacement (TMSD)^15^ and toehold exchange,^16^ illustrated in Fig. 1, are the key reactions underlying much of DNA nanotechnology. In these processes, an invading strand replaces another strand in a duplex by competing for base pairs. A double toehold exchange process provides a simple mechanism for DNA-mediated catalysis (Fig. 1 (b)); the invader *A* catalyses the replacement of *B* by *C* in the duplex with strand *D*. This simple mechanism also illustrates the subtlety of catalysis in general. Ideally, we would like each sub-process to be thermodynamically downhill so that reactions proceed in the desired direction – particularly if the reaction were part of a very complex network. We could enhance the overall drive (make Δ*G_AC_* more negative) by allowing *C* to bind to more bases of *D* than *B*. However, if these bases provide a toehold for *C* to initiate binding to the *BD* duplex, they will enhance the ability of *C* to displace *B* without the intervention of *A*, a leak reaction that compromises the catalytic control of *A* over the process.

**Figure 1:**
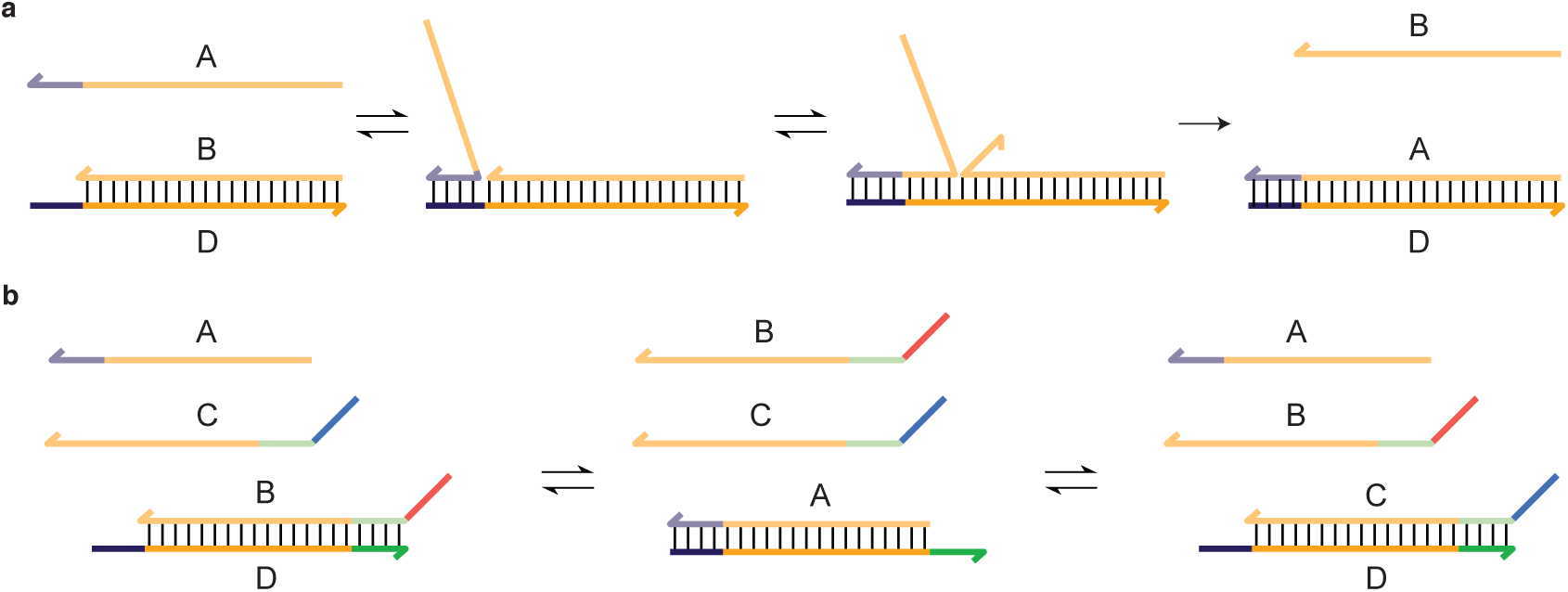
(a) Toehold-mediated strand displacement. The reactant duplex *BD* can be attacked by an invading strand *A* which is complementary to the whole of *D*, including an initially single-stranded toehold. (b) A simple catalytic motif engineered through toehold exchange. If displacement of *B* by *A* exposes a second toehold that allows the subsequent displacement of *A* by *C*, *A* can act as a catalyst for the interconversion of *BD* and *CD*, providing control over the reaction.

An ideal solution to this problem would be to provide a “hidden” thermodynamic advantageto the reaction, by which Δ*G_AC_* can be made substantially more negative without significiantly affecting the leak reaction rate. In this article, we propose the use of internal mismatches within the initial duplex (*BD* in Fig. 1 (b))) to fulfill this role. Through a combination of experiment and simulation, we demonstrate that, although an internal mismatch substantially destabilizes a duplex, its influence on the displacement rate is highly position-dependent: the mismatch location can be rationally chosen to provide a hidden thermodynamic drive.

Specifically, we first measure displacement rates for mismatch-repairing systems, finding that reaction rates are highest when the mismatch is repaired early but not immediately during strand-exchange, and that introduction of an initial mismatch can increase the reaction rate by two orders of magnitude whereas later mismatches have a very small effect. We use simulation to explain this behaviour, and confirm that later mismatches constitute a hidden thermodynamic drive. We then demonstrate rational design of hidden thermodynamic driving in a DNA pulse generator.

## 1 Results and Discussion

### 1.1 The dependence of displacement rate on mismatch position

Experiments to measure mismatch-repair TMSD reaction kinetics are shown in Fig. 2. Substrate duplexes *OT* contain a mismatch that is repaired by the invading strand *I* (Fig. 2). The final duplex *IT* thus has perfect complementarity. The toehold on the substrate duplex was chosen to be 4 nucleotides (nt) long and the displacement domain of *OT* was 20 base pairs (bp) in all cases. Single C-C mismatches were inserted in the displacement domain at positions between 2 and 17 bases from the toehold. To ensure the maximum degree of comparability between these systems, designs with mismatches at positions 2 → 13 were obtained by translating the mismatch sequence along the *OT* duplex, and mismatches at positions 15 and 17 were obtained by single point mutations to the target strand.

**Figure 2:**
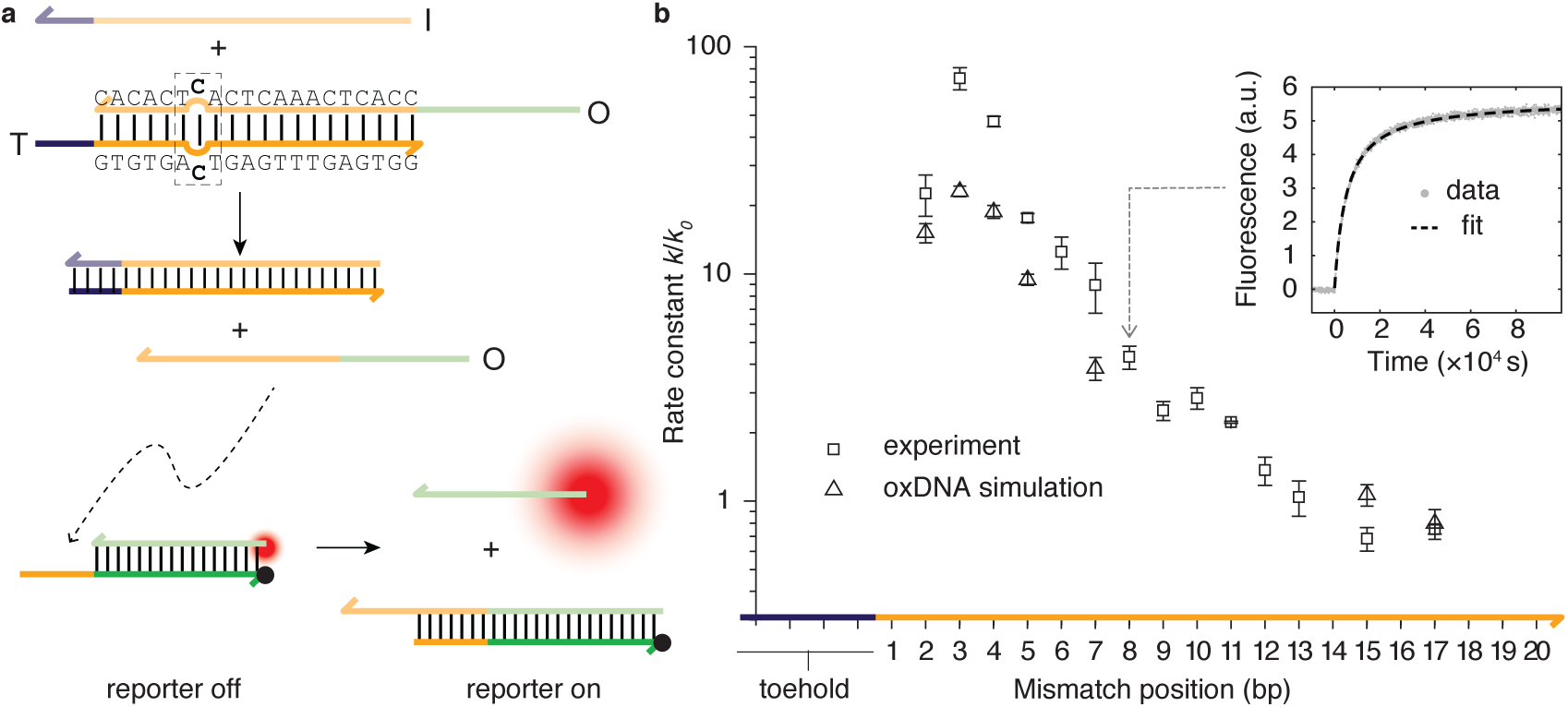
The effect of mismatch repair location on strand displacement rate. (a) Mismatch-repair TMSD. An invader *I* displaces an output strand *O* from an *OT* duplex containing a mismatch. *O* subsequently triggers a secondary displacement reaction that liberates a fluorophore-bearing strand from a quenching duplex. Complex *OT* has a cytosine-cytosine mismatch at position 7, counting from the beginning of the displacement domain. The 3-base pair mismatch motif (dashed rectangle) is the same for all mismatch positions. (b) Experimentally obtained TMSD rate constant as a function of mismatch position relative to *k*_0_ =2.4 × 10^3^ M^−1^s^−1^ for a mismatch-free system (squares). Fluorescence data and fit for position 7 is shown inset. Triangles show corresponding relative rate constants obtained from oxDNA simulations, showing a similar non-monotonic relationship between reaction rate and mismatch location. Error bars show SEM; there is also an error that systematically effects all relative rates due to the error in determination of no-mismatch rate constant *k*_0_ of ±0.14× for experimental points and ±0.11× for simulated points.

A fluorescent reporter reaction was used to measure displacement rates.^16–19^. Reporter fluorescence is triggered by displacement of single-stranded *O* from *OT*. Strand *O* possesses a5′ single-stranded domain that forms the displacement domain of the reporter interaction, but the toehold-binding domain for the reporter reaction is sequestered in the *OT* duplex until *O* is released. Liberated *O* displaces a fluorophore-labelled strand from the reporter duplex in which the fluorophore is quenched (Fig. 2). The reporter reaction is thus also based on the TMSD mechanism, but with a longer toehold of 7nt to ensure a significantly greater reaction rate. In this limit, the reporter provides a signal proportional to the cumulative output from *OT* at any given time (See Supplementary Information S1.1).

Fitted rate constants *k* for the mismatch-repair reaction varied between *k* ~ 1.8 × 10^5^ M^−1^ s^−1^ for position 3 and *k ~* 1.7 × 10^3^ M^−1^ s^−1^for position 15. Adjustment of the mismatch repair position can thus change the displacement rate by at least two orders of magnitude. The rate constant for an equivalent system with no mismatches is *k*_0_ =2.4 × 10^3^ M^−1^ s^−1^. Relative rates are plotted in Fig. 2 (b). The reaction rate constant rises as the mismatch is moved away from the toehold up to position 3; it then falls approximately exponentially back to the mismatch-free rate for positions ≳ 13. This nonmonotonicity is particularly striking when compare to the monotonic behaviour observed in a previous study of kinetic control through mismatch *creation*.^19^

### 1.2 Insights from coarse-grained simulations of DNA

The results in Fig. 2 suggest that mismatches far from any duplex ends could be used to provide ‘hidden’ thermodynamic driving: the effect of the introduction of the base pair mismatch on the reaction rate is limited despite a large predicted thermodynamic change of up to 6 kcal/Mol,^20^ approximately the contribution of three extra base pairs. To investigate this possibility, and to aid the rational design of such systems, we simulated mismatch repair displacement with a coarse-grained model, oxDNA.^21, 22^ oxDNA describes DNA as a string of rigid nucleotides with pairwise interactions that represent excluded-volume, backbone-connectivity, base-pairing and stacking interactions. It is simple enough that simulations of complex processes such as strand displacement are feasible whilst retaining enough of the underlying physics to capture the thermodynamic, mechanical and structural changes associated with the formation of duplex DNA from single strands. oxDNA has previously been applied to a number of systems in which displacement is important.^19, 23–25^. Here, we use the sequence-dependent parameterisation of Sulc *et. al*.^22^

Using oxDNA with forward flux sampling, as outlined in Section 3.6 and Supplementary Information S4.1 and S4.2, we obtained predictions for displacement rate constants for mismatch positions 2, 3, 4, 5, 7, 15 and 17 relative to the mismatch-free case (note that estimates of relative rates obtained from coarse-grained models are more accurate than absolute rates^23, 26^). Results are plotted in Fig. 2 (b), and a snapshot of the displacement process is shown in Fig. 3 (a). oxDNA reproduces the initial rise in rate up to mismatch position 3, followed by the decay back to the mismatch-free rate for late mismatch positions. The maximal acceleration, by a factor of 23, is comparable to, but smaller than, that observed in experiment. Much of this discrepancy may be due to known features of the oxDNA model: a slight enhancement of the tendency of short toeholds to mediate successful displacement^23, 25^ (limiting the potential speed-up due to mismatch repair), and a slight underestimation of the destabilisation of DNA due to mismatches.^21^

**Figure 3:**
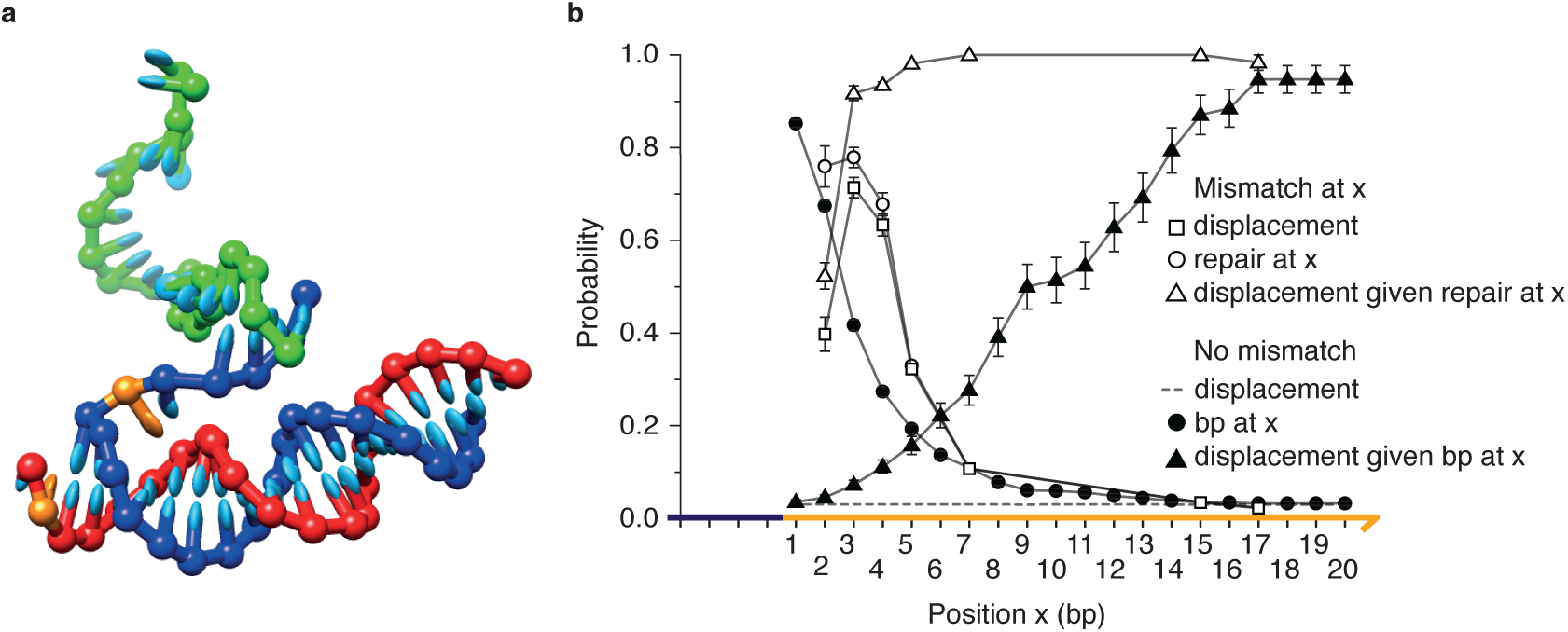
oxDNA simulation of mismatch repair TMSD. (a) Snapshot from an oxDNA simulation showing the invader (green) binding to the toehold of the target (blue); the output (red) is bound to the displacement domain. The mismatched base pair, highlighted in orange, is at position 2; fraying of the *OT* duplex is visible. (b) Analysis of strand displacement probabilities using oxDNA. The probability that the branch point reaches position *x* (*p*(*x* reached|toehold)) and the probability of successful displacement given that *x* is reached (*p*(disp| *x* reached)) are plotted for a perfectly-matched duplex (shaded symbols) and a duplex with a mismatches at *x* (open symbols). Also shown is the overall displacement probability *p*(disp|toehold) for the perfectly-matched case and *p*(disp|toehold) as a function of mismatch location. The presence of a mismatch at position *x* enhances both the probability of reaching *x* and of successful displacement given that *x* is reached.

More importantly, oxDNA elucidates the biophysical mechanism underlying the behaviour. In the second-order limit, relevant for weak, reversible toehold binding, the overall rate constant can be re-written as

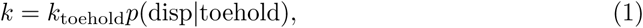

where *k*_toehold_ is the rate of toehold binding, and *p*(disp|toehold) is the probability of successful displacement (rather than abortive detachment) once the toehold duplex has formed. Given that the toehold is the same for all systems studied, *k*_toehold_ should be similar for all systems (as found in oxDNA simulations: see Supplementary Information S4.2). Therefore the key quantity is *p*(disp|toehold). Results for the first-order limit, which may be relevant at extremely high effective concentrations of reactants such as for surface-bound schemes ^27^, are presented in Supplementary Information S4.2.

We can split *p*(disp|toehold) further into

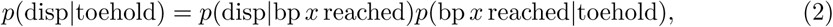

where *x* is any base-pair position in the displacement domain. The probabilities *p*(disp|bp *x* reached) and *p*(bp *x* reached|toehold) are plotted in Fig. 3 (b) for the mismatch-free system. Also shown are the probabilities for reaching position *x*, and the probability of success given that *x* has been reached, for a system with a mismatch at *x*. In the first case, the product is independent of the choice of reference base-pair *x*; in the second, it is strongly dependent on the position *x* of the defect. We see that the mismatch at *x* enhances both the probability that base pair *x* is reached and the probability of overall success given that *x* has been reached. Mismatches weaken the duplex *OT*, accelerating branch migration to the position of the defect, and discourage reverse branch migration once the repair is made. Both effects accelearate strand displacement. Both are suggested by the free-energy profile of displacement, which quantifies the stability of intermediate states as a function of reaction progression, plotted in Fig. 4. Lower free energies indicate more favourable states: it can be seen from Fig. 4 that the forward progress of displacement is encouraged by a large drop in free energy, and that this drop begins slightly before the mismatch is actually enclosed.

**Figure 4:**
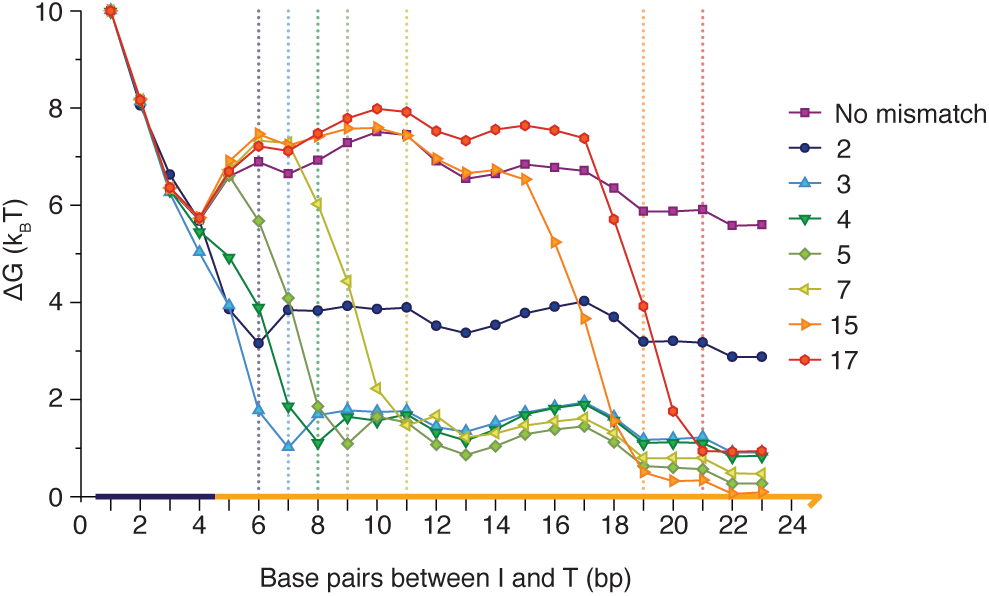
Free-energy profile of strand displacement as a function of branch point position. Simulated configurations are assigned to macrostates based on the number of base pairs between *I* and *T*. Mismatch repair is associated with a large drop in free energy which begins slightly before mismatch enclosure (dotted lines indicate the expected number of base pairs at the point of mismatch repair). A mismatch at position 2 provides an anomalously small overall thermodynamic bias for displacement. Error bars are not plotted (for clarity): overall free energy changes between 1 and 23 base pairs showed a typical standard error of *~*±0.2*k*_B_*T*

The magnitudes of these effects depend strongly upon *x*. For relatively late mismatches (*x* ≳7), the existence of the mismatch has almost no effect on the probability that this position is reached by the invader. The destabilising effect of the mismatch is quite local, affecting only nearby base pairs (Fig. 4): if the mismatch is sufficiently far from the toehold, altering the last few steps prior to the mismatch has almost no effect on the probability of reaching it. Additionally, once mismatches at large values of *x* are enclosed, the probability of eventual success is essentially unity. However, the probability of success having reached larger values of *x* is high even without the mismatch, so the overall increase in *p*(disp|toehold) due to the mismatch is quite small. The displacement rate therefore converges with the mismatch-free case for large *x*.

For early mismatches (small *x*) the presence of the defect significantly increases both *p*(disp|bp *x* reached) and *p*(bp *x* reached|toehold), suggesting that low values of *x* are optimal for accelerating displacement. However, *p*(disp|bp 2 reached) is much smaller than *p*(disp|bp 3 reached), leading to a substantial difference in rates and the non-monotonic dependence of displacement speed on mismatch position. The reason for this behaviour is that a mismatch at *x* = 2 is significantly less destabilising than a mismatch at *x* = 3.^21, 28^. Instead of fully enclosing the mismatch at position 2 the terminal base pair in the *OT* duplex can fray (Fig. 4), exchanging the high cost of incorporating a mismatch into the duplex for the lower cost of disrupting the terminal base pair. As a result, repairing the mismatch provides a smaller thermodynamic advantage for *IT* over *OT* than with a mismatch deeper inside the duplex, as can be seen by comparing the position-2 curve to the others in Fig. 4. A mismatch repaired at position 2 thus provides an anomalously small barrier to reversal of branch migration, hence *p*(disp| 2 reached) is relatively small.

The free energy profiles in Fig. 4, combined with the displacement rate data in Fig. 2 (b), indicate that mismatch repair can provide hidden thermodynamic driving when positioned appropriately. All repaired mismatches make a large, negative contribution to Δ*G* of reaction, but mismatches encountered later during branch migration have a very limited effect on reaction rates.

### 1.3 Demonstrating and exploiting hidden thermodynamic driving in a pulse-generating circuit

We now demonstrate our ability to design rationally a hidden thermodynamic drive by constructing a simple pulse-generating device in which two invaders compete to bind to a target using toeholds at opposite ends of the duplex. We label the strand that binds to the toehold at the 5′ end of the substrate duplex *I*_5_, and the strand that binds to the 3′ end *I*_3_ (as shown in Fig. 5 (a)). Each invading strand bears a quencher at its toehold end and a fluorophore at the other (fluorophore quencher pairs Cy3/IowaBlackFQ and Cy5/IowaBlackRQ for *I*_3_ and *I*_5_ respectively): binding to the target enhances the fluorescence from the successful invader as the formation of the target duplex separates the fluorophore from the quencher.

**Figure 5:**
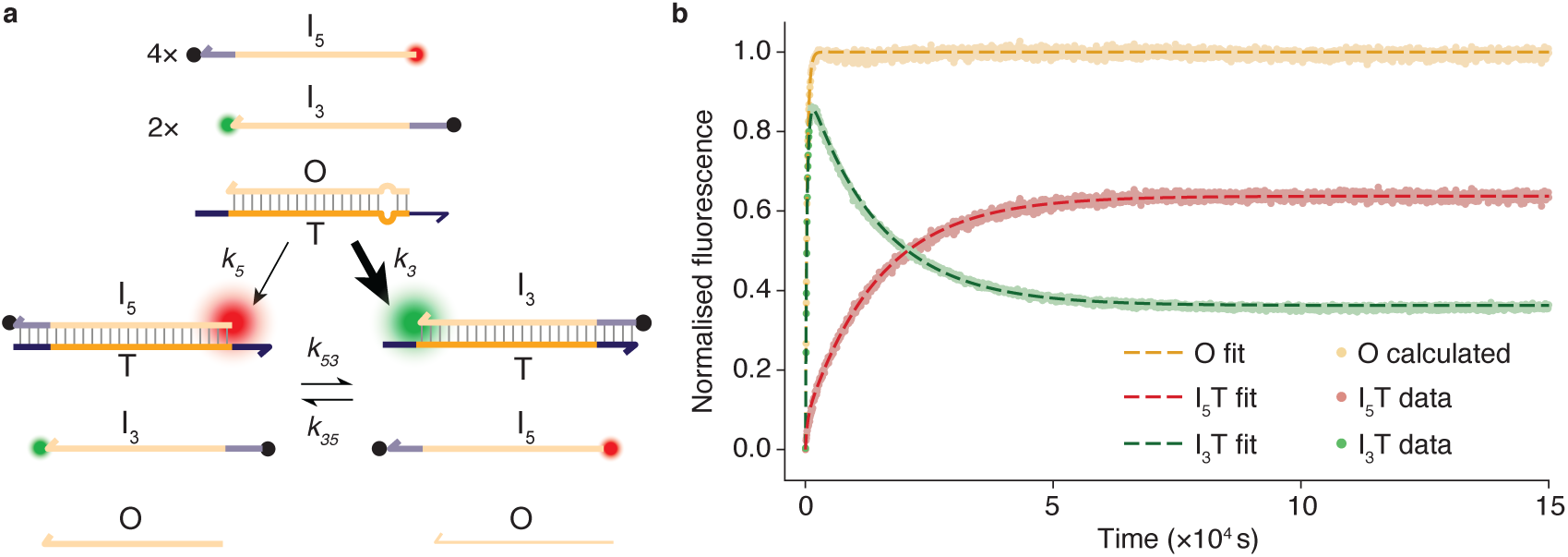
Pulse-generation with two-toehold competitive displacement reactions. (a) Schematic for two toehold competitive displacement reaction with an asymmetrically placed mismatch in the substrate duplex *OT*, 18 bases from the 5′ toehold and 3 bases from the 3′ toehold. The 3′ invading strand *I*_3_ should be kinetically favoured. 5′ invading strand *I*_5_ is thermodynamically favoured as it is present in greater excess than *I*_3_, and both repair the initially present mismatch. (b) T–C → T–A mismatch-repair at position 18 from the 5′ toehold of *OT*. Both processed data and ODE fits are shown (for fitted parameters see Supplementary Material S1.3). Initial relative concentrations are [*I*_5_]:[*I*_3_]:[*OT*] =4:2:1 initially.

The two toeholds are as similar as possible, but asymmetry is built into the system by incorporating a mismatch in the initial duplex at a position 18 base pairs from the 5′ toehold and 3 base pairs from the 3′ toehold. The mismatch is thus encountered early by the 3′ invader and late by the 5′ invader. We expect it to provide an additional thermodynamic advantage for both, but it should substantially speed up invasion by the 3′ invader only. If our hypothesis, that the slowdown of late mismatch repair relative to early mismatch repair is due to hidden thermodynamic driving rather than to a difference between the free energy changes in the two reactions, were true we would expect pulse-like behaviour upon simultaneous addition of *I*_5_ and *I*_3_ to a solution of *OT*. The kinetic advantage of *I*_3_ in displacing strand O is matched by a corresponding increase in the rate of the reverse reaction: the position of the equilibrium, which is determined only by the free energies of the competing reactions, is relatively insensitive to the position of the defect. The transient excess of *I*_3_ hybridized to T thus disappears as the system relaxes to equilibrium.

We challenge *OT* with a twofold excess of *I*_3_, the kinetically favoured invader, and a fourfold excess of *I*_5_, which shifts the eventual equilibrium towards *I*_5_*T* (Fig. 5 (b)). We indeed see an initial rapid rise in Cy3 fluorescence as *I*_3_ displaces strand *O* from the initial duplex *OT*, repairing a position-3 mismatch. Cy3 fluorescence reaches a sharp peak before falling again as *I*_3_ is itself displaced from the product duplex *I*_3_*T* by *I*_5_ via toehold exchange,^16^ reaching an equilibrium that is biased towards *I*_5_*T*. Fitting second order kinetics to the competing reactions provides support for the mechanism of hidden thermodynamic driving Positioning the mismatch 3 base pairs from the 3′ toehold increases the rate constant for invasion by *I*_3_ by a factor of approximately 20, whereas the *I*_5_ invader is accelerated by less than a factor of 2. The thermodynamic advantage of repairing the mismatch, which is enjoyed by both the *I*_5_ and *I*_3_ invaders, is effectively kinetically hidden for *I*_5_. Data fitting, further experiments with a mismatch close to the 5′ toehold, and a comparison to an alternative pulse-generating device that generates a less sharp signal,^19^ are discussed in Supplementary Material S1.3 and S3.1.

## 2 Conclusions

We propose that hidden thermodynamic driving – enhancing the free-energy change Δ*G* of a reaction whilst leaving reaction kinetics largely unchanged – is an important design principle for constructing synthetic non-equilibrium systems. Moreover, through a detailed analysis of mismatch repair during DNA strand displacement, we have demonstrated the possibility of engineering hidden thermodynamic driving in strand displacement networks.

We observe that although strand displacement rates are substantially enhanced when a mismatch is repaired early during displacement, by up to two orders of magnitude, the kinetic effect is minimal if the mismatch is repaired at a later stage. This behaviour, along with more subtle effects that lead to a non-monotonic dependence of displacement rates for early mismatches, is reproduced and rationalized through oxDNA simulations. These simulations also confirm that these results ar consistent with a largely position-independent contribution to Δ*G* of the mismatch, and therefore that the implementation of hidden thermodynamic driving is feasible.

We have studied 4-base toeholds, which also serve as a proxy for a ‘partially-matched’ longer toeholds, demonstrating our ability to enhance Δ*G* in such cases without substantially increasing leak reaction rates. The understanding gained from oxDNA simulations allows us to confidently predict behaviour for other toehold lengths. Length-zero toeholds are a particularly relevant limit, as this motif occurs frequently in the design of catalytically-controlled, kinetically-frustrated systems.^10,11,29^. In this case, both ends of the strand are equivalent, so mismatches at either end are effectively “early”, and significantly accelerate strand displacement. Centrally located mismatches in long displacement domains, however, will provide only weak enhancement of both conditional probabilities *p*(disp|bp *x* reached) and *p*(bp *x* reached|toehold), leading to an enhancement of displacement rates by only a factor of approximately 2 despite the large contribution to Δ*G*. We hypothesize that multiple mismatches can be included near the centre of a sufficiently long duplex, providing an even larger contribution to the free energy change associated with the reaction.

From a biophysical perspective, our results are consistent with previous studies on mismatch creation.^19^. However, the only previous attempt to systematically probe the consequences of mismatch repair did not see any signal of enhanced displacement, due to the use of much longer toeholds.^30^. Interestingly, we note that oxDNA predicts reaction acceleration even for longer toeholds if concentrations are high enough to access the first order limit (Supplementary Material S1.2). In this context, mismatch repair might be used to increase the fundamental speed limit of displacement reactions occuring, for example, at high effective concentrations between surface-localized reactants.^27^

Although previous work has also demonstrated the possibility of decoupling the thermodynamics and kinetics of strand displacement,^16, 19, 31^ the motifs therein do not allow for such a large *increase* in the overall thermodynamic drive whilst maintaining the extremely low direct reaction rates that are necessary for catalytic control. For example, enhancing Δ*G* by providing complementary bases in an externally accessible toehold will necessary enhance reaction rates substantially.

In principle, a hidden thermodynamic drive would allow us to push a reaction forwards thermodynamically without introducing longer toeholds that could significantly increase the rates of undesired reactions. The ability to do so would be extremely beneficial when considering the design of complex catalytic motifs in which certain reactions are deliberately kinetically frustrated. Similarly, in dynamic systems in which leaks due to partially-matched toeholds are a concern, it is beneficial to be able to enhance the Δ*G* of intended reactions without increasing toehold lengths and thereby encouraging spurious strand displacement.

**Acknowledgements:** NJH was supported by a DTC LSI grant, AJT by a Royal Society Wolfson Research Merit Award, and TEO by a Royal Society University Research Fellowship.

**Author Contributions:** NECH, TEO, JB and AJT conceived the project, following initial computational investigations by AG under the supervision of AAL and TEO. NECH, TEO, JB and AJT planned the experiments and simulations. NECH performed the experiments and analysis; TEO performed the simulations and analysis. NECH, TEO, JB and AJT wrote the paper.

## 3 Methods

### 3.1 Design rationale

#### 3.1.1 Mismatch type and location

The mismatch type used for all single-invader repair experiments was a cytosine-cytosine (C-C) mismatch, the most destabilising pairing according to the nearest-neighbour model as parameterised by SantaLucia.^28^. The invader strand *I* contains a guanine (G) at this position to form a correct Watson-Crick base pair and repair the C-C mismatch initially present. The *OT* duplex toehold is a 5′ single stranded overhang of the *T* strand; hence the invading strand *I* has its complement toehold on the 3′ end. For comparison with the two-toehold case we shall refer to this as 5′ invasion.

For all mismatch positions, the mismatch motif (the mismatch and its nearest neighbours) was preserved. For mismatch positions 1-13 this was achieved by sliding the mismatch domain along by moving the base pair immediately after it to the position immediately before it (so that all sequences are identical when the mismatch domain is omitted). This method also preserves base content and most of the nearest neighbour pairs and causes minimal change in free energy of reactant and product duplexes according to the nearest neighbour model.^28, 32^. Mismatch positions *>* 13 occur within the domain that doubles as the reporter toehold, therefore the output sequence must remain unchanged in this region. Mismatches can only be placed at locations (15 and 17) because they are cytosines in the output strand surrounded by the correct mismatch environment. By placing a cytosine where there was a guanine in the complementary domain the desired mismatch motif is achieved. Specific sequences are given in Supplementary Material S2.

#### 3.1.2 System design for two-toehold competition

For the two-toehold system an identical displacement domain sequence was used for the target strand *T* as in the single-invader experiment with the mismatch at position 3. Symmetric 4-nt sequences were added on either side of the displacement domain on *T* as toeholds. These toeholds have a different sequence to those used in the single-toehold experiments with a higher GC content. A strand *O*, complementary to the displacement domain of *T* except for a *C* − *T* mismatch at position 18, forms an initial duplex with *T*.

A5′ invader *I*_5_ has a 3′ domain complementary to the 5′ toehold of *T* to enable it to attack the two-toehold substrate duplex *OT* from its 5′ toehold. The 5′ invader was labelled with 5′ Cy5 and 3′ Iowa Black RQ with 2-nt spacers between the invader strand domains and either label. Similarly, the 3′ invader *I*_3_ bears the complement to the toehold domain at its 5′ end in order to attack the two-toehold duplex from its 3′ toehold. Labels on this invader were 5′ Iowa Black FQ and 3′ Cy3 with the same spacer nucleotides before the labels. The fluorescence change observed upon displacement comes from an increase in fluorophore-quencher separation when the invader strand is part of a duplex with the target. Sequences for all strands invovled in the two-toehold experiment are provided in Supplementary Material S2.

### 3.2 Secondary reporter systems

Two different reporter systems were used to generate the results presented in this paper. The first, Reporter A, had a 7-nt toehold domain as a single-stranded 5′ overhang and a 16-nt displacement domain. An additional 2 bp were present at the end of the reporter duplex opposite to the toehold. These last two base pairs are not directly invaded by the output, but will rapidly dissociate spontaneously after branch migration.^16, 19^. The output strand *O* is intended to trigger the reporter. Initially, the requisite toehold domain of *O* is sequestered within the *OT* duplex and the displacement domain exists as a single-stranded 5′ overhang.

In preliminary studies it was discovered that Reporter A is subject to leak reactions when the mismatches to be repaired are in the part of the O*T duple*x that sequesters the reporter toehold domain (positions 15 and 17). In this case the substrate duplex can displace the reporter output at a significant rate even in the absence of invader I, presumably due to spontaneous fraying which reveals the toehold of O. To avoid this effect an alternative reporter *B wa*s designed that extended the displacement domain of Reporter A by 2 base pairs into the toehold domain, with a compensating 2-nt 5′ extension of the toehold. The result is that the 7-nt toehold for Reporter B is more effectively sequestered within the O*T duplex*, starting 2 nt from the end of the duplex. Unlike Reporter A, Reporter B cannot be used with mismatches in position 12 or 13 since the toehold of *O fo*r triggering the reporter would be disturbed in these cases. Sequences, and evidence that the two reporter designs were consistent, are provided in Supplementary Material S1 and S2.

### 3.3 Oligonucleotide and complex preparation

Oligonucleotides were purchased from Integrated DNA Technologies. Strands without modifications were purchased with standard desalting; no additional purification was performed. Strands with fluorophore and/or quencher modifications were HPLC-purified by IDT. Strands were lyphophilised by IDT and resuspended to 100 μM in ddH_2_O for storage. Single-stranded species were diluted to 10 μM in fluorometry buffer FB (10 mM Tris·HCl pH 8.0 + 50 mM NaCl + 10 mM MgCl_2_). Reporter complexes were produced by incubating reporter output and reporter target at room temperature in FB at a ratio of 6:5, determined by titration. The nominal concentration of the reporter complex is the nominal concentration of reporter target used to produce it, usually 20 nM. Output:Target substrate duplexes were made by mixing components 1:1 in FB to a concentration of 10 nM and annealed by heating to 96◦C and cooling to room temperature at a rate of 6◦Cmin^−1^.

### 3.4 Spectrofluorometry

Fluorescence measurements were carried out in a JY Horiba Spex Fluoromax-3 spectrofluorometer with 4-position sample changer. Excitation wavelength was 648±2 nm, emission wavelength was either 664±1 nm, 664±2 nm or 664±5 nm. Integration time for measurements was 2 s and the interval between successive measurements was 60 s. Samples were not illuminated between measurements to minimise photobleaching. Fluorometer temperature was held at 25◦C with a water bath.

1.92 ml samples were added to a set of 4 matched quartz cuvettes (Hellma) with approximate volume of 2 ml and sealed tightly with 1.5 ml Eppendorf tubes as lids to reduce photobleaching and evaporation effects by decreasing contact of sample solution with air. 2.4 μl of 20 μM reporter complex was added to 1.92 ml buffer and a fluorescence baseline recorded. After baseline collection 2.4 μl of 10 μM output-target duplex was added and the sample mixed for approximately 60 s before returning to the fluorometer and collecting a second baseline. This enabled verification that there is no significant leak reaction between unreacted output-target duplex and reporter complex. After collection of a second baseline 2.4 μl of 10 μM invader strands were added and mixed, marking t=0 and the experiment start point.

For two-toehold experiments, invading strands were added to 1.92 ml FB to achieve the desired concentrations and a baseline was collected in the fluorometer. Of the 4 sample positions in any fluorometer run, three were used for experiments and one as a control. In the three experiment cuvettes, 2.4 μl of 10 μM *OT* duplex was added and mixed with the solution of invaders at t=0. Concurrently the fourth cuvette with the same concentration of invaders was saturated with a 10× excess of two-toehold target strand *T* to provide a measure of the maximum achievable fluorescence and to provide a control for global fluorescence fluctuations.

After use, cuvettes were washed 2× with 100% ethanol then 5× with ddH2O and finally 2× with ethanol again before drying with hot air and polishing the exterior with ethanol-soaked lens tissue.

### 3.5 Data processing and fitting

Data processing and fitting was carried out using MATLAB 2014b. Fluorescence traces were processed and fitted to ordinary differential equation models based on ideal second order kinetics. Details are provided in Supplementary Material S3.

### 3.6 oxDNA methods

oxDNA simulations were performed using the sequence-dependent version of the model presented by Sulc *et. al*.^22^ at a temperature of *T* = 25◦C. All simulations were performed on a single set of *O, T, I* strands in a periodic box of volume 16686 nm^3^. Sequences corresponded to the experimental design.

Kinetic studies were performed using the “Langevin B” algorithm for rigid bodies of Davidchack *et al*., ^33^ with friction parameters set to γ =0.586 ps^−1^ and Γ=1.76 ps^−1^ and using a time step of 8.53 fs.^23, 24^ Forward Flux Sampling^34, 35^ was used to enhance sampling of the dynamics of the displacement process, with individual dynamical trajectories used to infer the conditional probabilities reported in Fig. 3 (b). Details of the FFS, and the statistics of sampling runs, are reported in Supplementary Material S4. Relative fluxes for different systems give a good estimate of the relative rate constants obtained in the dilute second-order limit, as discussed in Ref.^19, 23, 36^

Thermodynamic sampling, used to obtain the free-energy profile and overall free energies of reactions, was performed using the Virtual-Move Monte Carlo method of Whitelam and Geissler^37, 38^ (specifically, the variant in the appendix of Ref.^38^). Seed moves were the rotation of the a nucleotide about its backbone site (angle drawn from a Gaussian distribution with standard deviation 0.12 radians) and translation of a nucleotide (distance drawn from a Gaussian distribution with standard deviation 0.102 nm). Umbrella Sampling^39^ was employed to enhance sampling efficiency. Details of the umbrella sampling procedure are provided in Supplementary Material S4.

